# Sleep Deprivation Alters Hippocampal Dendritic Spines in a Contextual Fear Memory Engram

**DOI:** 10.1101/2025.03.02.641043

**Authors:** Matthew Tennin, Hunter T. Matkins, Lindsay Rexrode, Ratna Bollavarapu, Samuel D. Asplund, Tanya Pareek, Daniel Kroeger, Harry Pantazopoulos, Barbara Gisabella

**Affiliations:** Department of Psychiatry and Human Behavior, University of Mississippi Medical Center, Jackson, MS, USA; Program in Neuroscience, University of Mississippi Medical Center, Jackson, MS, USA; Department of Anatomy, Physiology, and Pharmacology, College of Veterinary Medicine, Auburn University

**Author notes:** **Address for correspondence:** Barbara Gisabella, PhD, Department of Psychiatry and Human Behavior University of Mississippi Medical School, TRC 408, 2500 North State St Jackson, MS 39232, USA.

## Abstract

Sleep is critically involved in strengthening memories. However, our understanding of the morphological changes underlying this process is still emerging. Recent studies suggest that specific subsets of dendritic spines are strengthened during sleep in specific neurons involved in recent learning. Contextual memories associated with traumatic experiences are involved in post-traumatic stress disorder (PTSD) and represent recent learning that may be strengthened during sleep. We tested the hypothesis that dendritic spines encoding contextual fear memories are selectively strengthened during sleep. Furthermore, we tested how sleep deprivation after initial fear learning impacts dendritic spines following re-exposure to fear conditioning. We used ArcCreER^T2^ mice to visualize neurons that encode contextual fear learning (Arc+ neurons), and concomitantly labeled neurons that did not encode contextual fear learning (Arc-neurons). Dendritic branches of Arc+ and Arc-neurons were sampled using confocal imaging to assess spine densities using three-dimensional image analysis from either sleep deprived (SD) or control mice allowed to sleep normally. Mushroom spines in Arc+ branches displayed decreased density in SD mice, indicating upscaling of mushroom spines during sleep following fear learning. In comparison, no changes were observed in dendritic spines from Arc-branches. When animals were re-exposed to contextual fear conditioning 4 weeks later, we observed lower density of mushroom spines in both Arc+ and Arc-branches, as well as lower density of thin spines in Arc-branches in mice that were SD following the initial fear conditioning trial. Our findings indicate that sleep strengthens dendritic spines in neurons that recently encoded fear memory, and sleep deprivation following initial fear learning impairs dendritic spine strengthening initially and following later re-exposure. SD following a traumatic experience thus may be a viable strategy in weakening the strength of contextual memories associated with trauma and PTSD.

## Introduction

The idea that memories are strengthened or consolidated during sleep is well-established (for review see ^1^). However, evidence regarding the morphological and molecular underpinnings of memory consolidation during sleep has only recently emerged. A series of studies by Tononi and Cirelli developed the synaptic homeostasis hypothesis of sleep ^2,3^. This hypothesis posits that as organisms interact with their environment and encode new memories during wakefulness, neurons form new synapses and strengthen existing ones. During sleep, when the active encoding process is offline, synapses are downscaled to prevent over-excitation of neurons and improve signal-to-noise ratio and memory performance ^2,3^. In support of this hypothesis, several studies have reported an overall decrease in dendritic spines, synaptic density, and synaptic markers during sleep in sensory and motor cortical regions ^4–6^. However, an opposing view suggests that rather than collectively downscaling all synaptic connections, specific synapses that were recently involved in memory formation may be strengthened during sleep ^7–9^.

Several recent studies also suggest that the process of dendritic spine regulation during sleep is not a uniform downscaling of all synapses, but a more complex process including selective upscaling and downscaling of specific dendritic spines depending on recent experiences ^10–13^. For example, Born and colleagues propose that selective synapses formed during the day are tagged by specific proteins for strengthening during sleep to reorganize memory storage ^14,15^. Support for this theory comes from data demonstrating that sleep deprivation (SD) impairs memory strength ^7^, and results in decreased dendritic spines ^8^, suggesting that selective synapses may indeed be strengthened during sleep. For example, dendritic spine formation in specific branches of layer V motor cortex neurons is increased after motor learning ^9^, even though cortical areas may be subjected to an overall net synaptic downscaling during sleep ^4,5^. Short-term (5 hours) of SD was shown to result in reduced long term plasticity in CA1 hippocampal neurons ^7^, and reduced dendritic spine density ^8^, suggesting sleep is required for synapse stabilization in this region.

What may be the cause of the discrepancies in theories regarding dendritic spine changes during sleep? One critical factor in answering this question is dissociating which neurons were recently active and engaged in learning and which were not. For example, contextual memories associated with stressful experiences such as fear are necessary for survival and represent recent learning in dendritic spines that may be strengthened during sleep. By tagging dendritic spines of neurons recently engaged in fear learning, we can separate them from others and determine whether either population experiences up- or downscaling.

To determine the effect of sleep on neurons recently involved in learning, we used ArcCreER^T2^ mice, which allowed us to permanently label fear memory engram neurons ^16–19^. By concurrently labeling non-engram neurons using a viral labeling approach, we created two sets of clearly delineated neuronal populations which allowed us to assess whether dendritic spines that encode contextual fear memory are upscaled during sleep. This is highly relevant since several recent studies suggest that SD following emotional learning may be a promising preventative measure for post-traumatic stress disorder (PTSD)^20–22^.

Furthermore, since repeated re-exposure to trauma is often a significant contributor to PTSD risk and symptom severity ^23–27^, we examined the effect of sleep on contextual fear memory strength following a second exposure to a traumatic event. Specifically, we tested the hypothesis that SD following initial fear learning not only disrupts dendritic spine upscaling following the first exposure but may also disrupt this process following re-exposure to contextual fear conditioning 4 weeks later.

## Methods

### Animals

We used adult (3-4 months old) male ArcCreER^T2^ mice (Jackson Labs stock# 022357). Animals were maintained on a 12:12 h light-dark cycle with ad libitum access to water and food. All animal procedures met National Institutes of Health standards, as outlined in the Guide for the Care and Use of Laboratory Animals, and all protocols were approved by the University of Mississippi Medical Center Institutional Animal Care and Use Committee (IACUC). These mice express Cre-ER^T2^ under the direction of an Activity Regulated cytoskeletal associated protein (Arc) promoter region that is activated during fear-conditioned learning ^17–19,28^. These mice have been well characterized for selectivity of Arc expression in the fear memory engrams ^17–19,28^. The Arc protein is localized to dendritic spines, particularly in sites of increased postsynaptic activity ^29^. Importantly, extensive evidence supports the role of Arc in memory formation and consolidation (for review see ^29^ and ^30^).

### Viral Vector Injection

ArcCreER^T2^mice were anesthetized with 1.5% isoflurane and received bilateral stereotaxic microinjections of a cocktail of 2 μl of AAV mCherry ChR2 viral vector under the control of the CMV promoter (AAV2-CMV-DIO-mCherry), and 2 μl of AAV1-CMV-eGFP virus (Vector Biolabs, cat# 7002) in each hemisphere. The virus cocktail was infused using a Nanofil 35-gauge stainless steel bevel needle (catalog # NF35BV, World Precision Instruments, Inc., Sarasota, FL) attached to a 10μl Nanofil syringe (Hamilton Company, Reno, NV) as described in our previous study ^31^. Hamilton syringes were mounted in a stereotaxic barrel holder, and the rate of virus delivery was controlled by an automated syringe pump (Harvard Apparatus, Holliston, MA). The injections targeted the CA1 region of the dorsal hippocampus (stereotaxic coordinates AP: −2.1, ML:1.75, DV:1.0). The AAV2 serotype was selected for the mCherry ChR2 viral vector as this serotype has been reported to result in minimal spread and shows preference for labeling neurons over other cell types ^32^ (Vector Labs, cat# 7104). Specifically, injections of the AAV virus under the CMV promoter allowed for the labeling of dendritic branches for spine analysis. The neurons participating in the memory trace are labeled because the contemporaneous injection of 4-OH-tamoxifen allows the CreER to enter the cell nucleus, where it may cause recombination events at loxP-flanked sites, permanently marking the cells that participated in that memory trace. Cells that are transduced with viral vectors allow for Cre-dependent expression of mCherry, permanently marking the cells that participated in that memory trace in red. This design allowed for visualization of neurons that are part of the fear memory engram, expressing both mCherry for Arc and YFP (Arc+). Neurons not part of the fear memory engram (Arc-) express only GFP. After 6 weeks, animals were divided randomly into two cohorts of four mice each (4 controls, and 4 SD mice).

### Contextual Fear Conditioning

Mice were given 36 days for recovery from surgery and optimal viral vector expression. Then they were placed in darkness for 12 hours to avoid non-specific Arc-YFP labeling as established by previous studies ^16–19^, (Fig. 1A). Mice were injected with 4-OH-tamoxifen in the middle part of their prior dark cycle (1 AM). 4-OH-tamoxifen in mice has been established to reach detectable levels rapidly, peak between 5-9 hours following injection, and disappear almost completely by 36 hours, thus restricting stimuli via dark housing during this time window reduces non-specific YFP expression ^17,28,33,34^. Four hours later (5 AM), the mice underwent auditory fear conditioning to activate CreERT2 production. Fear conditioning was conducted as in our previous study ^35^. This time point is chosen to allow fear-conditioned animals to sleep during their normal sleep period after fear learning. Animals were either allowed to sleep normally afterwards or underwent 5 hours of SD using an automated sleep deprivation procedure (see below). Animals were then placed back into a dark environment for 48 hours following fear conditioning to avoid non-specific Arc-YFP labeling, as established by previous studies ^17–19,28^. We used two experimental groups of 4 control and 4 SD animals in each group. The first group was used to examine the effect of SD on dendritic spines immediately following fear learning (Fig. 1A). The second group of mice was used to examine the effect of SD following initial fear learning on dendritic spines after re-exposure to the fearful stimulus 4 weeks later (Fig. 1B). Our experimental timelines were guided by similar procedures reported in prior studies using ArcCreER^T2^ (Cazzulino et al. 2016)

**Figure 1.**
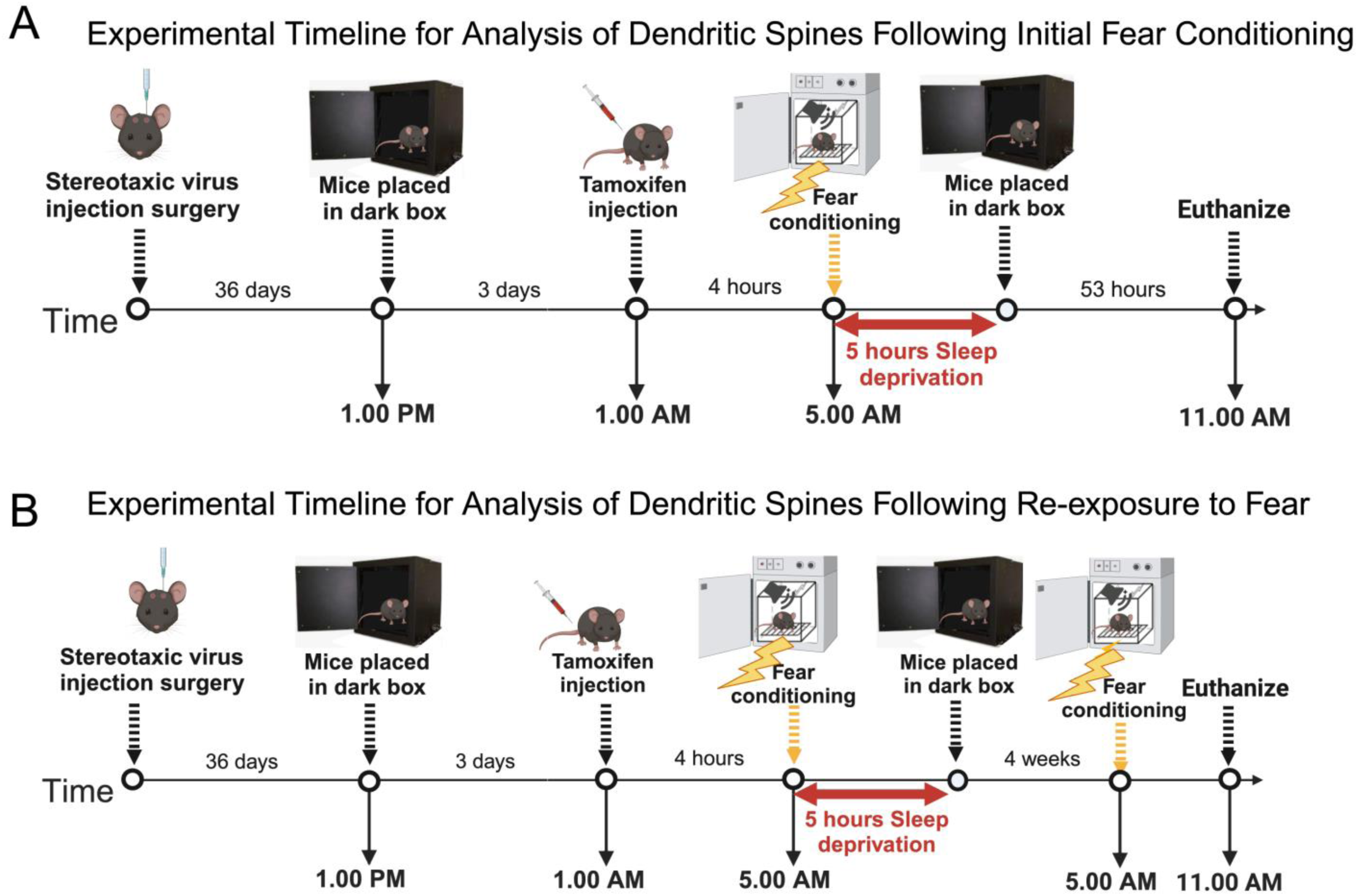
Experimental timeline. <u>Sleep Deprivation and Dendritic Spine Analysis After Initial Fear</u> <u>Learning:</u> ArcCreER^T2^ mice were stereotaxically injected with dual AAV viral vectors to label dendritic spines involved in contextual fear learning (mCherry, Arc positive, Arc+) and spines not involved in contextual fear learning (eGFP, Arc negative, Arc-). (A) After 36 days, mice were placed in darkness for 12 hours to avoid any non-specific Arc induction, injected with 4-OH-tamoxifen at 1 AM, and 4 hours later underwent contextual fear conditioning, followed by 5 hours of sleep deprivation or normal sleep. Animals were then placed back into a dark environment for 48 hours and euthanized for brain sample processing. (B) <u>Dendritic Spine Analysis After Contextual Fear Re-exposure:</u> In the second set of experiments, animals underwent a similar timeline of procedures to label dendritic spines followed by fear conditioning with or without SD. These animals were then returned to standard housing for 4 weeks before undergoing re-exposure to contextual fear conditioning. Mice were allowed to sleep for 5 hours and were then sacrificed for brain sample processing.

### Automated Sleep Deprivation

The Pinnacle automated sleep deprivation system (Cat# 9000-K5-S) which simulates gentle handling was used for 5 hours of sleep deprivation from lights on (6 AM) to 11 PM. This system consists of a cylindrical housing chamber with a bar that continuously rotates at 5 rpm and randomly reverses direction every 10–30 s, which prevents animals from sleeping. All animals were housed in the cylindrical chambers beforehand to adapt to the environment and control animals were housed in the same chambers but without the rotating bar. A researcher visually verified that the bar always rotated and that mice did not use alternative strategies to avoid the bar and sleep. This system has been established by prior studies to effectively reduce sleep as measured with EEG recordings in rats and mice ^31,36–39^. Both control and SD mice were sacrificed at 11 AM and processed for quantification of dendritic spines (Fig. 1). This time point (5 h into the light cycle) is identical to previous studies from our lab and others ^31,40,41^. Brains were then analyzed for viral vector expression and dendritic spine density as in our published studies ^31,40^.

### Tissue processing

Mice were sacrificed by cervical dislocation and perfused with 0.1M phosphate-buffered saline (PBS) containing 4% paraformaldehyde. Brains were removed and cryoprotected in 30% sucrose in 0.1M PBS (pH 7.4), then sectioned into coronal 40 μm serial sections using a freezing microtome (American Optical 860, Buffalo, NY) and stored in 0.1M phosphate buffer with 0.1% NaAzide. Sections were mounted on gelatin-coated slides and coverslipped using Dako mounting media (S3023, Dako, North America, Carpinteria, CA) to quantify dendritic spine density from images captured using confocal microscopy.

### Confocal imaging

A Zeiss LSM 880 confocal microscope system interfaced with Zen imaging software (ZEN 2.3 SP1) was used to acquire 3D image stacks of dendritic branches from hippocampal CA1 neurons in sections from control and SD mice. All slides used for confocal imaging were coded for blind analysis. Images of 40 μm-thick sections were acquired, with a z-step of 0.5 μm using a 63x oil immersion objective (numerical aperture 1.4 DIC M27; pixel size, 0.10 x 0.10 μm) similar to the method described in our previous study ^31,35^. For dendritic spine quantification, confocal microscopy images were analyzed using Neurolucida 360 software with Autospine to measure spine density in hippocampus CA1 cells using an approach previously described ^35,40,42,43^. Dendritic spines were sampled in the CA1 area of the hippocampus by an investigator who was blinded to the treatment group. We collected confocal images of all visible viral vector-labeled dendritic branches in stratum radiatum (apical dendrites) within the CA1 area of the hippocampus using 25×25 μm confocal imaging windows. Analysis was restricted to stratum radiatum, as this was the area where the majority of Arc+ dendrites were observed. Slides were coded for blind analysis, and sampling boxes were placed in the CA1 area. The length from the border of CA2 to the medial end of CA1 was measured for each section, and sampling boxes were placed on either side of the mid-point between these two ends of CA, halfway between the lateral border of CA1/CA2 and the medial end of CA1. Two sampling boxes (450 x 300 μm^2^) were placed in the stratum radiatum each 300 μm left and right of the midline, positioned immediately below the pyramidal cell layer. We then imaged all dendrites located in each box visible with Arc+ (mCherry red), or Arc-(YFP green) fluorescence.

### Dendritic spine quantification

For dendritic spine quantification, confocal microscopy images were analyzed using Neurolucida 360 software with Autospine to measure spine density as described in our previous study ^31,35,40^. Spines were grouped into thin spines, stubby spines, and mushroom spines automatically by the Neurolucida 360 software based on the spine head to neck diameter ratio (1.1), length-to-head ratio (2.5) mushroom head size (0.35 μm), and filopodia length (3 μm) according to previously established criteria ^43^. Spine density, shape, and volume were quantified using Neurolucida 360 with semiautomated analysis from 3D confocal image stacks in an unbiased manner.

### Statistical analysis

Densities of dendritic spines were calculated as spines per dendrite segment length in micrometers. For all statistical tests, the significance threshold was p ≤ .05. Non-parametric (Wilcoxon-Mann-Whitney) tests were used to compare population estimates for control and SD groups as data were not normally distributed and were followed by Bonferroni post hoc correction. Box plots were used to depict the data for each group from n= 4 control and n= 4 SD animals).

## RESULTS

### Sleep deprivation following fear learning decreases dendritic spines in the contextual fear memory engram

Arc+ branches were largely distributed in stratum radiatum of CA1 (Fig 2A), and dendritic spine densities were quantified from Arc+ and Arc-branches (Fig. 2B,C). No significant difference was observed in overall spine density between SD and control mice (Fig. 2 D). We observed significantly lower density of mushroom spines in SD mice compared to control mice (p<0.004, Fig. 2E). Overall densities of thin and stubby spines did not differ between SD and control animals (Fig. 2E). Decreased densities of mushroom spines were observed in Arc+ branches of SD mice, indicating that the upscaling of mushroom spines during sleep following fear learning is driven by neurons that encoded the recent contextual fear memory (p<0.001, Fig. 2G). In comparison, densities of stubby spines were increased in Arc+ branches of SD mice in comparison to controls (p<0.05, Fig. 2G). No changes in dendritic spine densities from Arc-branches were detected between SD and control animals (Fig. 2H,I).

**Figure 2.**
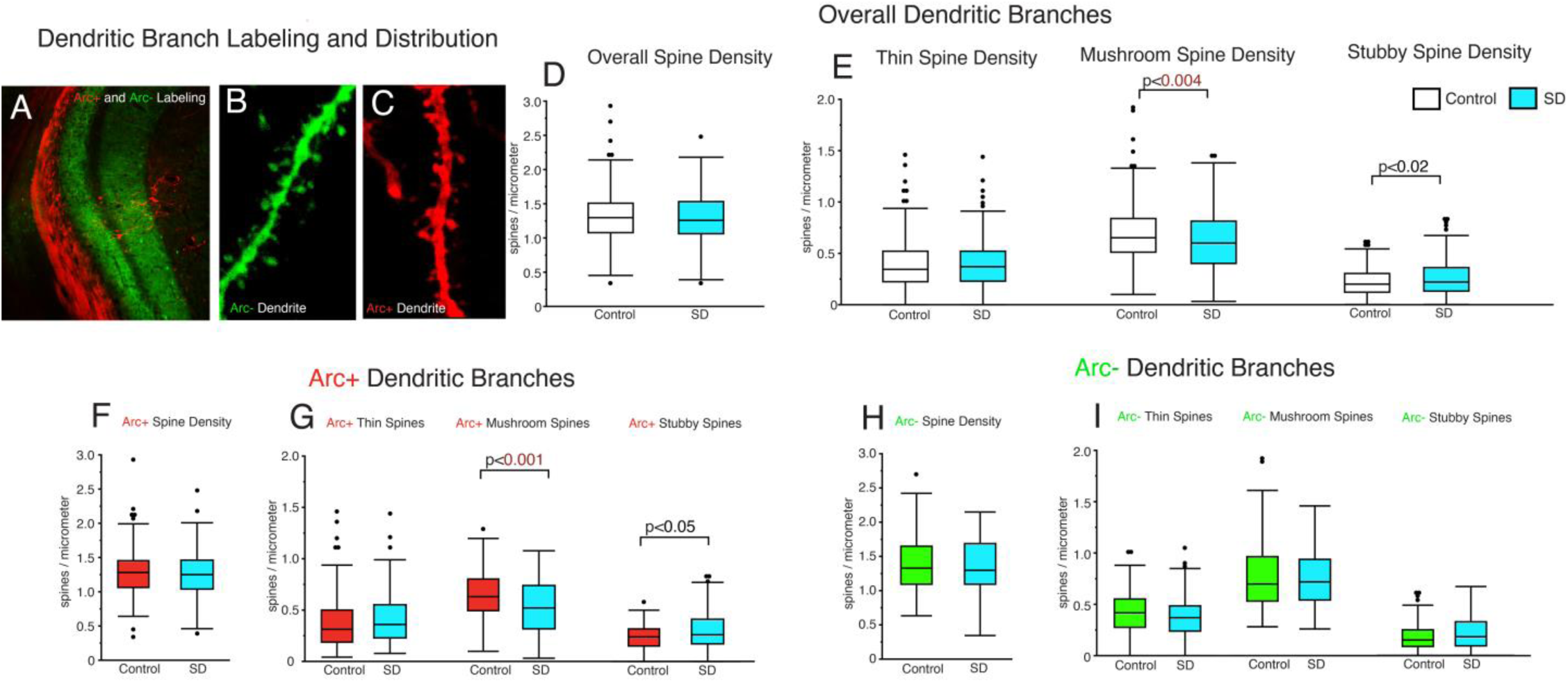
Sleep deprivation following initial fear learning decreases dendritic spines in the contextual fear memory engram. The majority of Arc+ branches in CA1 were distributed in stratum oriens (A). Confocal images depicting representative branches and spines from Arc-(B) and Arc+ (C) neurons in CA1. No difference was observed in overall spine density between control and SD mice (D). When comparing overall spine densities for specific spine classes, there was significantly lower density of mushroom spines in SD mice (n_mice_ = 4; n_dendrites_ = 223) compared to control mice (n_mice_ = 4; n_dendrites_ = 260) indicating synaptic upscaling during sleep (E). No differences were detected for densities of thin or stubby spines. No difference in overall Arc+ spine density was detected between SD and control mice (F). Arc+ branches displayed decreased density of mushroom spines in SD mice (n_mice_ = 4; n_dendrites_ = 135) compared to control mice (n_mice_ = 4; n_dendrites_ = 110), indicating upscaling of mushroom spines during sleep in the recently encoded contextual fear memory engram. In contrast, we observed an increase in stubby spine density in Arc+ branches, and no changes were observed in thin spines (G). In comparison, Arc-branches displayed no differences in dendritic spine density (SD: n_mice_ = 4; n_dendrites_ = 88; control: n_mice_ = 4; n_dendrites_ = 114)(H-I). Box plots depict values for each group, statistical significance was determined using the Wilcoxon-Mann-Whitney test.

### Sleep deprivation following fear learning alters dendritic spine head and neck length

Dendritic spine neck length is associated with the rate of synaptic transmission and synaptic strength, with shorter length associated with faster synaptic transmission ^44^. Significantly shorter spine head and neck length was observed specifically for thin spines (head length: p<0.002; Fig. 3A and neck length: p<0.001; Fig. 3B) from Arc+ branches in SD mice compared to control mice. No changes in spine head length were detected in mushroom and stubby spines in Arc+ branches (Fig. 3A,B). In comparison, Arc-branches displayed significantly shorter spine head length in mushroom spines (head length: p<0.005) and stubby spines (p<0.02; Fig. 3C), along with greater head length of thin spines (p<0.005; Fig. 3C) in SD mice compared to control mice. Neck backbone length in spines from Arc-branches was increased in thin spines (p<0.01); Fig. 3D) and decreased in stubby spines (p<0.001; Fig. 3D) from SD mice compared to control mice.

**Figure 3.**
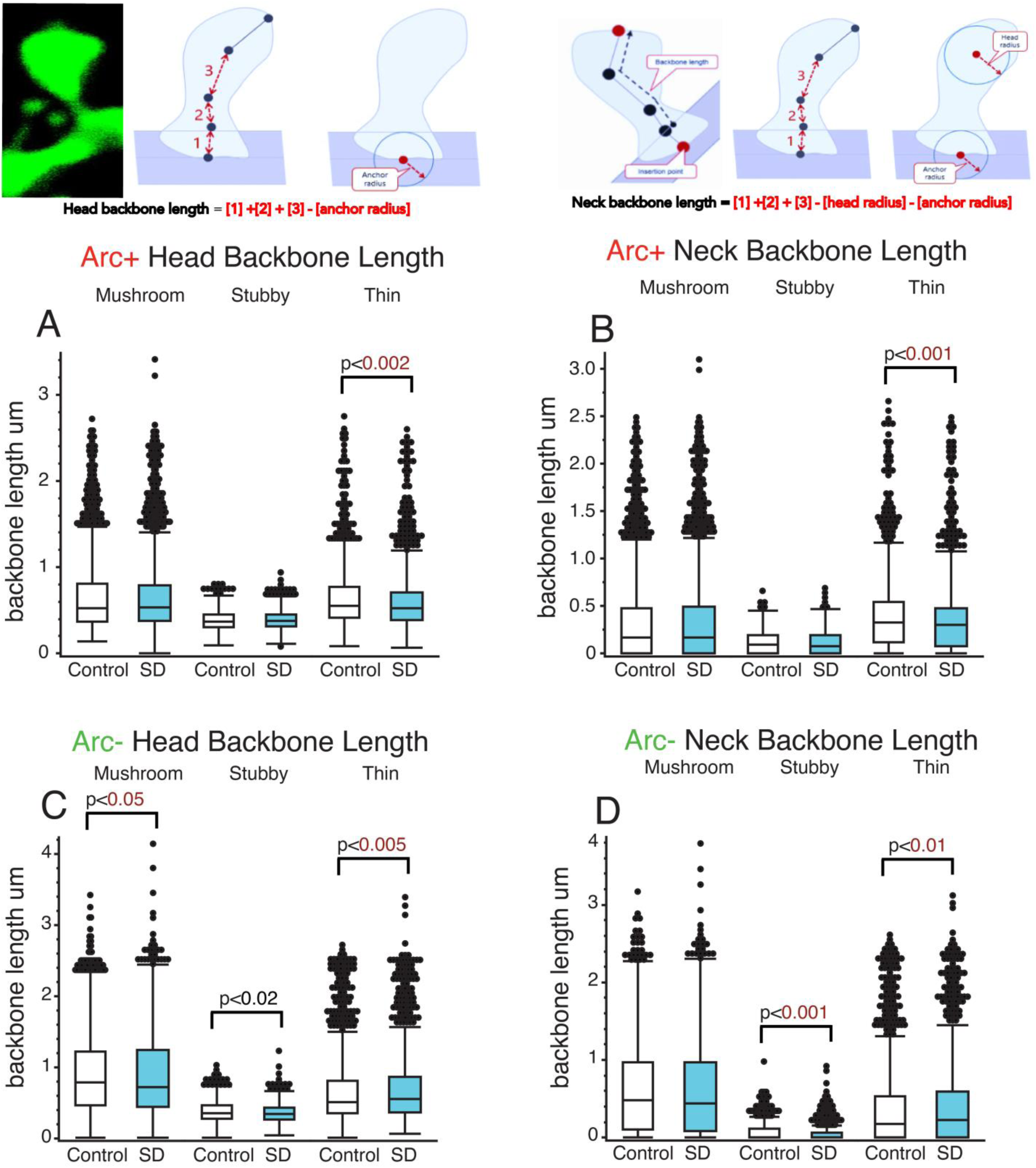
Sleep deprivation following fear learning alters dendritic spine head and neck length. The diagram depicts neck backbone length measured as the distance from the insertion point to the center of the spine head minus the head radius and anchor radius. Head backbone length is measured as the distance from the insertion point to the center of the spine head minus the anchor radius. (A,B) Significantly shorter head and neck length of thin spines was observed in Arc+ branches from SD mice compared to control mice (control; n_mice_ = 4; n_spines_ = 2289; SD; n_mice_ = 4; n_spines_ = 2063). No differences were observed in Arc+ branches for head and neck length of mushroom (control; n_mice_ = 4; n_spines_ = 1938; SD; n_mice_ = 4; n_spines_ = 2966) or stubby spines (control; n_mice_ = 4; n_spines_ = 818; SD; n_mice_ = 4; n_spines_ = 1444). Arc-branches displayed significantly shorter spine head length in mushroom spines (control; n_mice_ = 4; n_spines_ = 2717; SD; n_mice_ = 4; n_spines_ = 2055) and stubby spines (control; n_mice_ = 4; n_spines_ = 759; SD; n_mice_ = 4; n_spines_ = 661) in SD vs control mice (C,D). Greater head length of thin spines was also observed in Arc-branches of SD mice (C) spines (control; n_mice_ = 4; n_spines_ = 1563; SD; n_mice_ = 4; n_spines_ = 1061). Box plots depict values for each group, statistical significance was determined using the Wilcoxon-Mann-Whitney test.

### Sleep deprivation after fear learning alters dendritic spine head and neck diameter, volume, and surface area

Dendritic spine neck diameter is normally positively correlated with head volume ^45,46^. No changes in overall spine head and neck diameter were detected in thin, mushroom, or stubby spines of Arc+ branches in SD mice compared with control mice after fear conditioning (Fig. 4A-B). In contrast, Arc-branches from SD mice displayed greater head and neck diameter specifically in stubby spines (head diameter: p<0.001; Fig. 4C; neck diameter: p<0.001; Fig. 4D). Spine volume and surface area is a measure of synaptic strength and is positively associated with the density of glutamate receptors ^47–50^. Smaller spine surface area and volume in Arc+ branches was observed in thin spines from SD mice versus control mice (volume: p<0.03; Fig. 4E; surface area: p<0.03; Fig. 4F). Moreover, we observed greater surface and volume for stubby spines in Arc-branches (volume : p<0.03; Fig. 4G; surface area: p<0.01; Fig. 4H).

**Figure 4.**
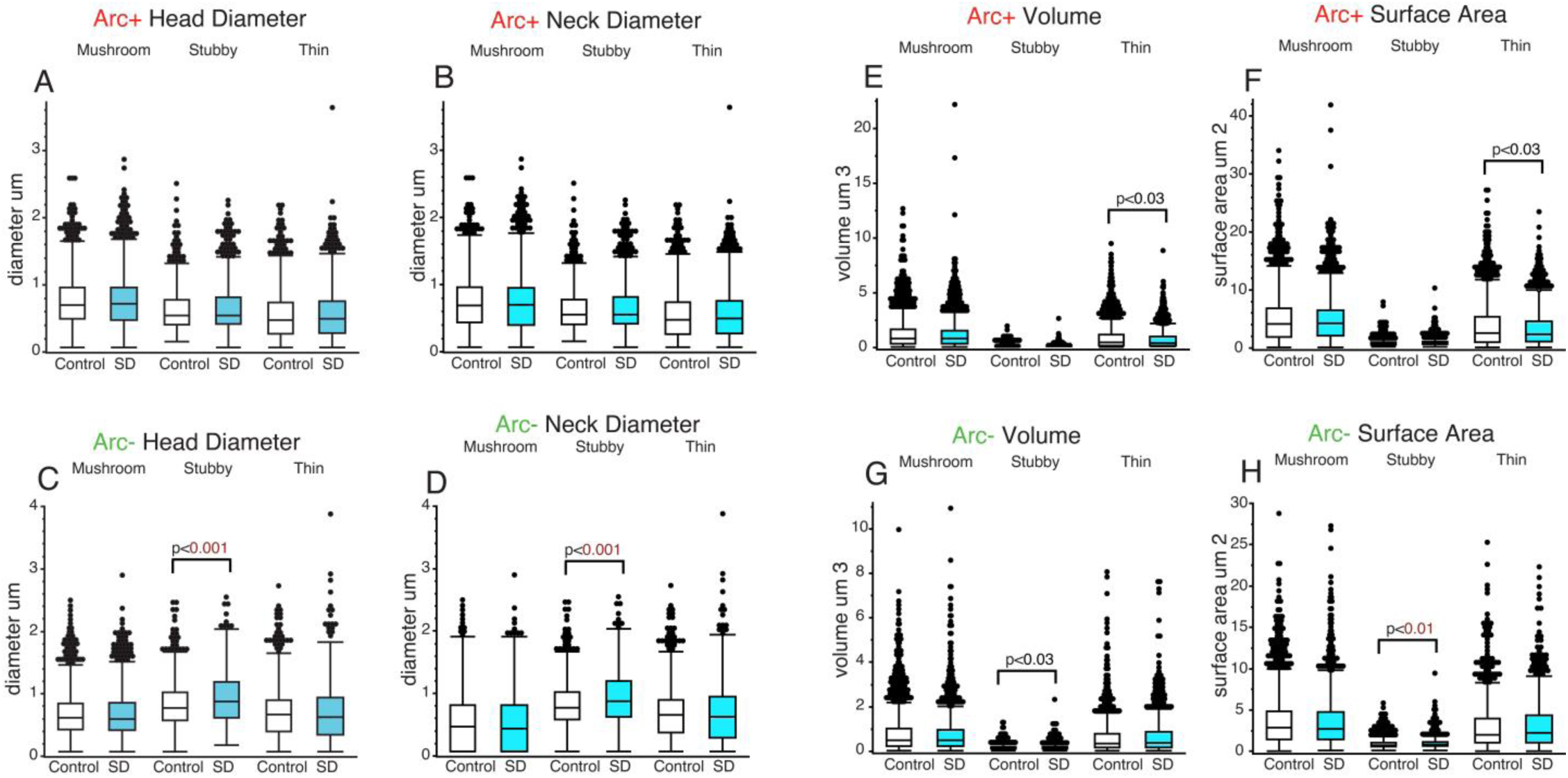
Sleep deprivation after fear learning alters spine head and neck diameter, volume and surface area. No changes in head and neck diameter were detected in mushroom spines (control; n_mice_ = 4; n_spines_ = 1938; SD; n_mice_ = 4; n_spines_ = 2966), stubby spines (control; n_mice_ = 4; n_spines_ = 818; SD; n_mice_ = 4; n_spines_ = 1444), or thin spines (control; n_mice_ = 4; n_spines_ = 2289; SD; n_mice_ = 4; n_spines_ = 2063) spines from Arc+ branches in SD mice comparted to control mice (A). Arc-branches displayed greater head and neck diameter of stubby spines in SD mice compared to control mice (control; n_mice_ = 4; n_spines_ = 759; SD; n_mice_ = 4; n_spines_ = 661), (C,D). Smaller volume and surface area was observed in thin spines from Arc+ branches in SD mice compared to control mice (control; n_mice_ = 4; n_spines_ = 1563; SD; n_mice_ = 4; n_spines_ = 1061), (E,F). Greater surface area and volume was observed for stubby spines from Arc-branches in SD mice compared to control mice (E,F). Box plots depict values for each group, statistical significance was determined using the Wilcoxon-Mann-Whitney test.

### Sleep deprivation following initial fear learning disrupts dendritic spine upscaling that occurs after re-exposure 28 days later

We tested the hypothesis that SD after the initial fear conditioning session weakens synaptic upscaling that occurs following future re-exposure to the traumatic experience by re-exposing mice to fear conditioning 28 days later. Spine density across all spines (Arc+ and Arc-combined) was significantly decreased in SD mice compared with control mice after fear conditioning re-exposure (p<0.001; Fig. 5A), indicating synaptic upscaling during sleep following re-exposure. This decrease was selective for thin spines (p<0.007; Fig. 5B). Mice that were sleep disrupted following the initial fear conditioning session displayed a trend towards lower freezing during re-exposure 28 days later, with significantly lower freezing duration during the first post-tone interval and increasing to control levels by the 3^rd^ post-tone interval (Fig. 5C). Moderate effects were observed in dendritic spine densities from Arc+ branches. No change was observed in overall density of Arc+ spines (Fig. 5D), possibly due to opposite changes in mushroom vs thin spines. Mushroom spine density was significantly decreased in SD mice compared to control mice (p<0.001; Fig. 5E), whereas thin spine density was increased in SD mice (p<0.002; Fig. 5E). In comparison, Arc-branches displayed an overall decrease of dendritic spine density in SD mice (p<0.001; Fig. 5F), that was significant for thin (p<0.001; Fig. 5G) and mushroom spines (p<0.003; Fig. 5G). Stubby spines from Arc-branches displayed significantly greater density in SD mice (p<0.02; Fig. 5G).

**Figure 5.**
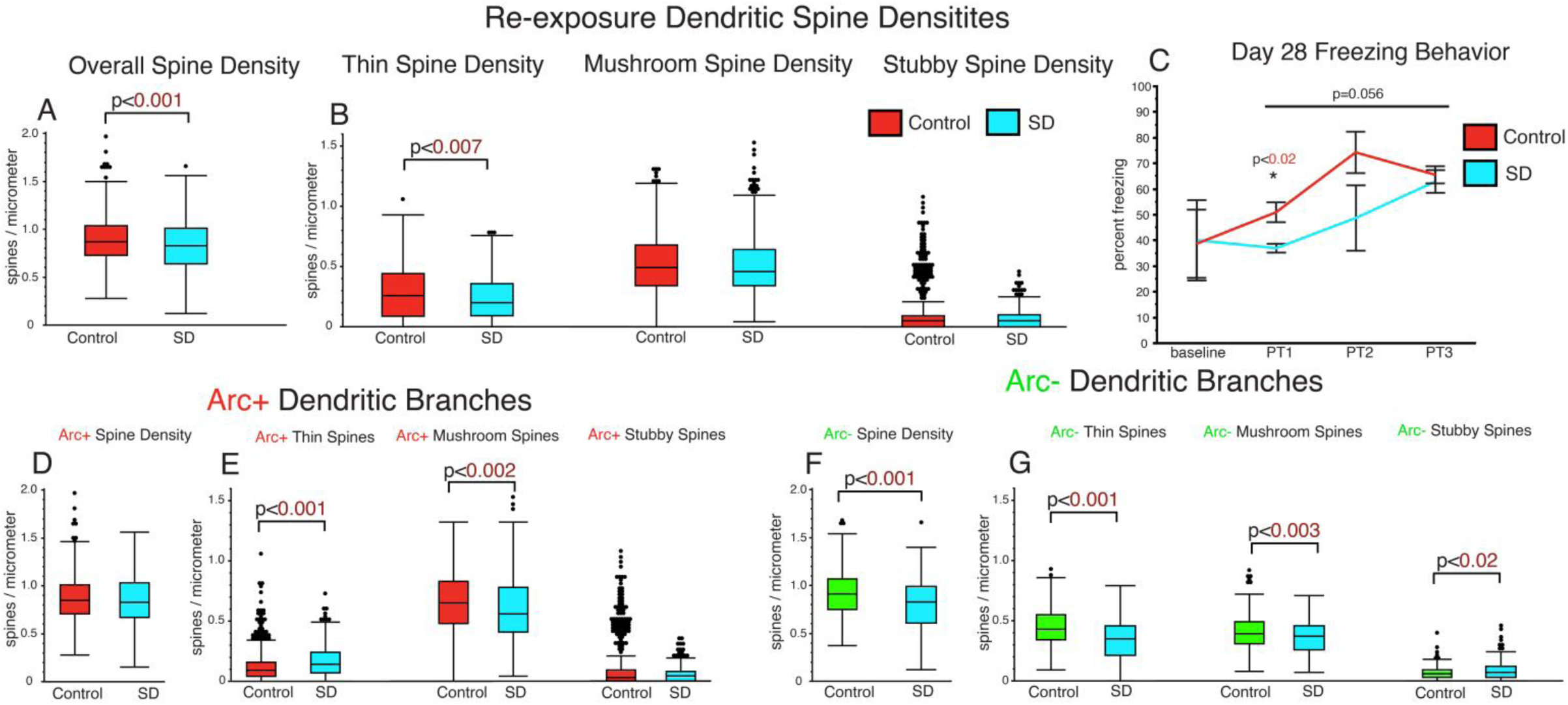
Sleep deprivation following initial fear learning disrupts dendritic spine upscaling that occurs after re-exposure 28 days later. The overall spine density across both Arc+ and Arc-spines was significantly lower in SD mice (n_mice_ = 4; n_dendrites_ = 768) versus control mice (n_mice_ = 4; n_dendrites_ = 1245) following re-exposure to fear conditioning 28 days after the initial exposure (A). This decrease was selective for thin spines (B). Mice that were sleep disrupted following initial exposure to fear conditioning displayed a trend towards less freezing during re-exposure, with significantly lower freezing during the first post-tone (PT) interval (C). No difference was observed in overall spine density from Arc+ branches (D). Mushroom spine density was significantly lower in Arc+ branches from SD mice (n_mice_ = 4; n_dendrites_ = 484) compared to control mice compared to control mice (n_mice_ = 4; n_dendrites_ = 452) (E). Thin spine density was increased in SD mice compared to control mice (E). Arc-branches displayed overall lower dendritic spine density in SD mice (n_mice_ = 4; n_dendrites_ = 284), compared to control mice (n_mice_ = 4; n_dendrites_ = 571)(F). Lower density was present in thin and mushroom spines (G). Stubby spines from Arc-branches displayed significantly greater density in SD mice (G). Box plots depict values for each group, statistical significance was determined using the Wilcoxon-Mann-Whitney test. Plots for fear conditioning data represent means and 95% confidence intervals and significance was determined using ANOVA for individual time points and repeated measures ANOVA across all timepoints.

### Sleep deprivation following initial fear learning alters dendritic spine head and neck length after re-exposure to fear conditioning 28 days later

Thin spines from Arc+ branches from SD mice displayed significantly shorter head and neck length compared to control mice (head length: p<0.001; Fig. 6A and neck length: p<0.001; Fig. 6B) following re-exposure to fear conditioning. Greater neck length was observed for mushroom spines from Arc+ branches in SD mice compared to control mice (neck length: p<0.04; Fig. 6B). No difference was observed for head or neck length in stubby spines from Arc+ branches between SD and control mice (Fig. 6A,B). In comparison Arc-branches of SD mice displayed greater head and neck length of thin spines (head length: p<0.001; neck length: p<0.001), (Fig. 6C-D) and greater head length for mushroom spines (head length: p<0.04). Shorter head and neck length was observed for stubby spines from Arc-branches of SD mice compared to control mice (head length: p<0.02; and (p<0.001; Fig. 6C-D).

**Figure 6.**
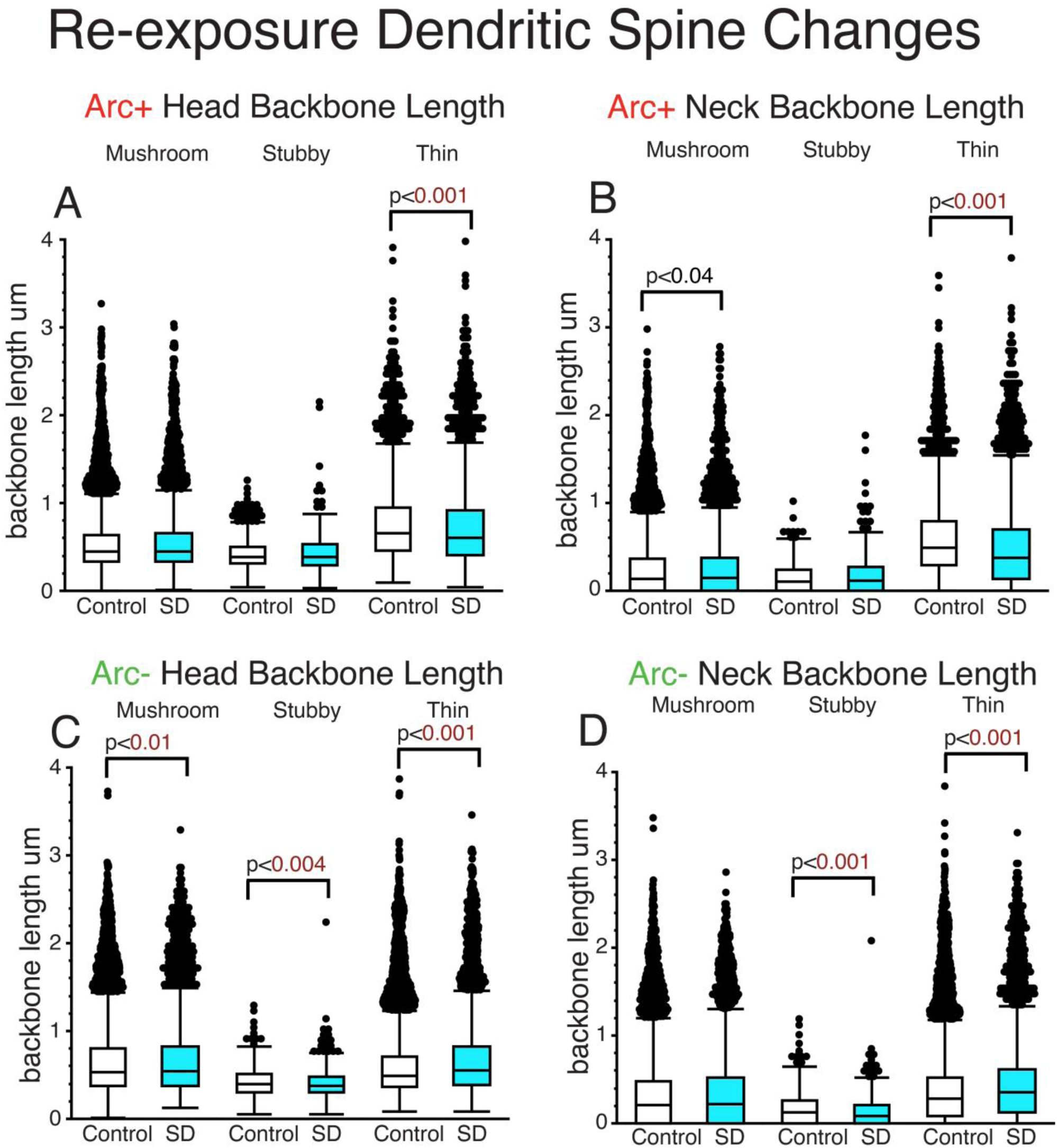
Sleep deprivation following initial fear learning alters dendritic spine head and neck length after re-exposure to fear conditioning. Thin spines from Arc+ branches from SD mice displayed significantly shorter head and neck length compared to control mice (control; n_mice_ = 4; n_spines_ = 2654; SD; n_mice_ = 4; n_spines_ = 2403) following re-exposure to fear conditioning (A,B). Greater neck length was observed for mushroom spines from Arc+ branches in SD mice compared to control mice (control; n_mice_ = 4; n_spines_ = 12375; SD; n_mice_ = 4; n_spines_ = 7894), (B). No difference in head and neck length was observed in stubby spines from SD mice compared to control mice (control; n_mice_ = 4; n_spines_ = 2106; SD; n_mice_ = 4; n_spines_ = 901), (A,B). Arc-branches displayed greater head and neck length of thin spines (control; n_mice_ = 4; n_spines_ = 10642; SD; n_mice_ = 4; n_spines_ = 3871) and head length of mushroom spines (control; n_mice_ = 4; n_spines_ = 9129; SD; n_mice_ = 4; n_spines_ = 4208) in SD mice compared to control mice (C,D). Shorter head and neck length was observed for stubby spines from Arc-branches of SD mice compared to control mice (control; n_mice_ = 4; n_spines_ = 1540; SD; n_mice_ = 4; n_spines_ = 957), (C,D). Box plots depict values for each group, statistical significance was determined using the Wilcoxon-Mann-Whitney test.

### Sleep deprivation following initial fear learning alters dendritic spine head and neck diameter and spine size after re-exposure to fear conditioning 28 days later

Smaller head diameter was observed for mushroom spines from Arc+ branches from SD mice compared to control mice (p<0.06; Fig. 7A), along with greater head and neck diameter of thin spines (head diameter: p<0.001; Fig. 7A; neck diameter: p<0.001; Fig. 7B). In comparison, greater head diameter was observed in mushroom spines from Arc-branches (head diameter: p<0.04; Fig. 7C), along with greater head and neck diameter of stubby spines (head diameter: p<0.001; Fig 7C; neck diameter: p<0.001; Fig. 7D) in SD vs control mice. Thin spines from Arc-branches displayed smaller neck diameter in SD mice (neck diameter: p<0.001; Fig. 7D). Smaller volume of mushroom spines was detected in Arc+ branches from SD mice (p<0.001; Fig. 7E), along with smaller volume and surface area of stubby spines (volume: p<0.001; surface area: p<0.001; Fig. 7E&F). Increased volume and surface area were detected for all spine types from Arc-branches from SD mice compared to control mice (mushroom: volume: p<0.001; Fig. 7G; surface area: p<0.001; Fig. 7H; stubby: volume: p<0.001; Fig. 7G; surface area: p<0.001; Fig. 7H; thin: volume: p<0.001; Fig. 7G; surface area: p<0.001; Fig. 7H).

**Figure 7.**
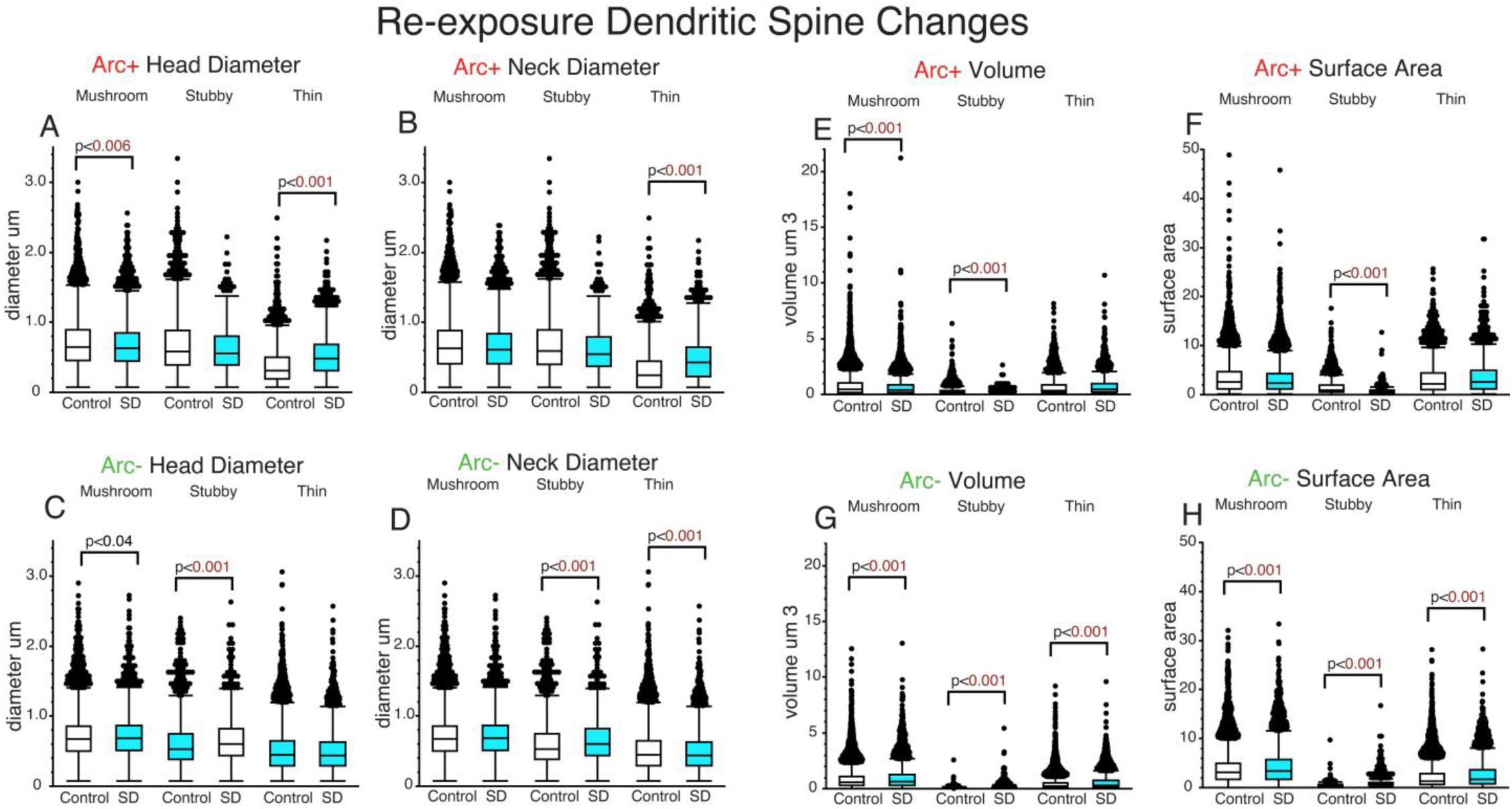
Sleep deprivation following initial fear learning alters dendritic spine head and neck diameter and spine size after re-exposure to fear conditioning. Smaller head diameter was observed for mushroom spines from Arc+ branches from SD mice compared to control mice (control; n_mice_ = 4; n_spines_ = 12375; SD; n_mice_ = 4; n_spines_ = 7894) following re-exposure to fear conditioning 28 days after initial exposure (A). Greater head and neck diameter of thin spines was observed in Arc+ branches from SD mice compared to control mice (control; n_mice_ = 4; n_spines_ = 2654; SD; n_mice_ = 4; n_spines_ = 2403), (A,B). Arc-branches displayed greater head diameter of mushroom spines (control; n_mice_ = 4; n_spines_ = 9129; SD; n_mice_ = 4; n_spines_ = 4208) and greater head and neck diameter of stubby spines (control; n_mice_ = 4; n_spines_ = 1540; SD; n_mice_ = 4; n_spines_ = 957) in SD mice compared with control mice (C,D). Thin spines from Arc-branches displayed smaller neck diameter in SD mice compared with control (control; n_mice_ = 4; n_spines_ = 10642; SD; n_mice_ = 4; n_spines_ = 3871), (D). Smaller volume of mushroom spines was detected in Arc+ branches from SD mice compared with control mice, along with smaller volume and surface area of stubby spines (E,F). Increased volume and surface area were detected for thin, mushroom and stubby spines from Arc-branches from SD mice compared to control mice (G,H). Box plots depict values for each group, statistical significance was determined using the Wilcoxon-Mann-Whitney test.

## DISCUSSION

Our results demonstrate that SD following contextual fear learning disrupts upscaling of dendritic spines in the fear memory engram of CA1 neurons (Fig. 8). This represents, to our knowledge, the first evidence for synaptic upscaling during sleep in a hippocampal fear memory engram and provides key insight regarding reports of synaptic upscaling vs synaptic downscaling in this region during sleep ^8,40,51^. Furthermore, our findings indicate that re-exposure to a traumatic event results in upscaling during sleep in neurons that were not part of the original contextual memory engram, thus expanding the engram, and that SD after initial fear learning impairs this synaptic upscaling that occurs following re-exposure (Fig. 5). Furthermore, mice that were sleep disrupted following the initial fear conditioning session displayed a trend towards less freezing during re-exposure to fear conditioning 28 days later (Fig. 5). Together our results provide insight into the contextual fear engram specific morphological alterations that occur during sleep and support the potential therapeutic and protective effect of SD following a traumatic experience.

**Figure 8.**
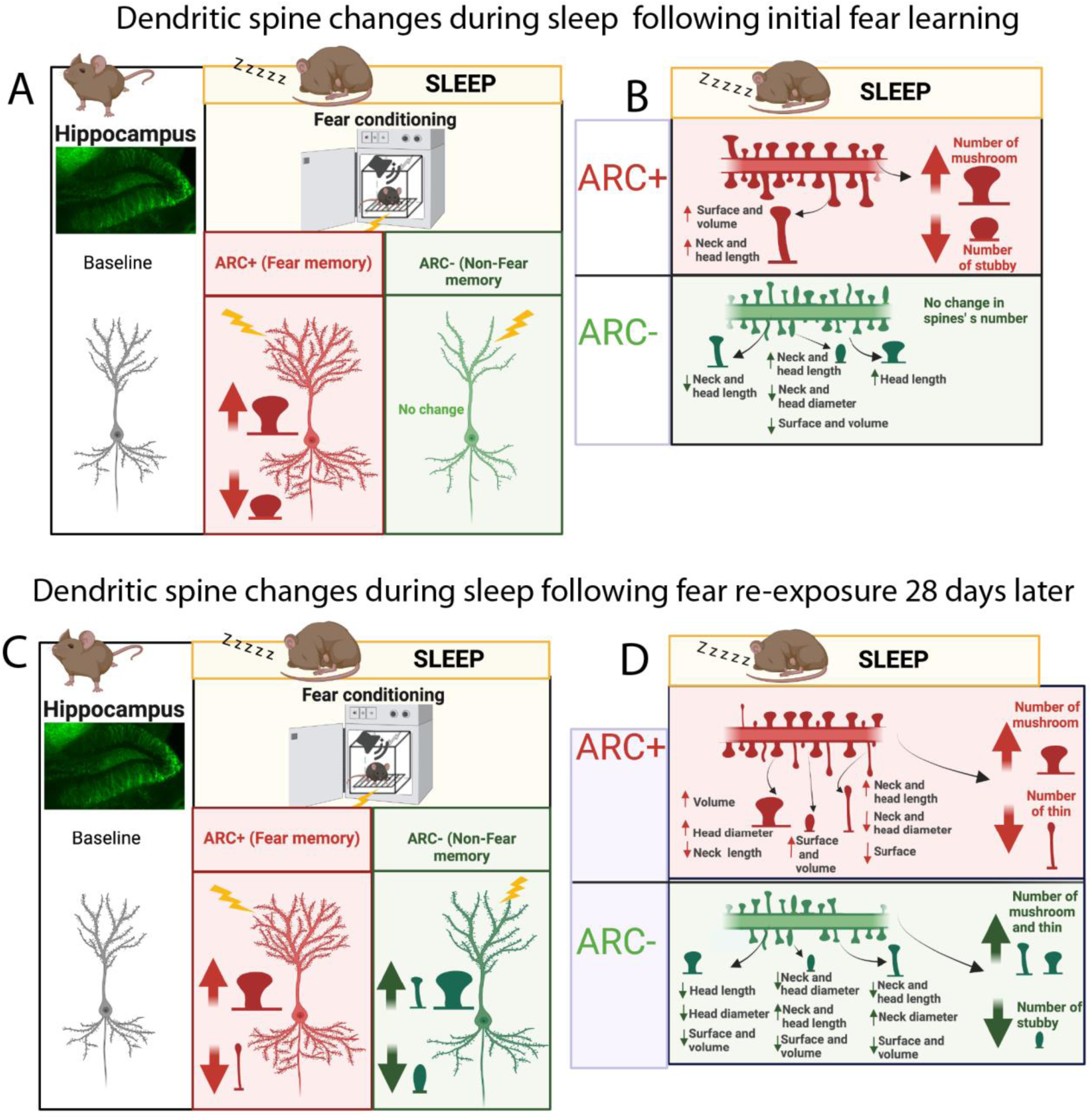
Summary of Dendritic Spine Changes During Sleep in the CA1 Contextual Fear Memory Engram. The diagram represents a summary of the working hypothesis of dendritic spine changes during sleep in CA1 neurons that initially encoded fear memory (Arc+) and neurons that did not (Arc-). Spine densities for each region are indicated by the branch with multiple spines in the upper part of each panel, and spine morphological subtypes are indicated by corresponding shapes (thin, subby, mushroom) in each panel. (A) Densities of mushroom spines are increased in Arc+ neurons during sleep following initial contextual fear learning. (B) Surface area, volume head, and neck length of Arc+ thin spines are increased during sleep, whereas head and neck length of mushroom spines is increased in Arc-branches. (C) Densities of Arc+ mushroom spines and Arc-thin and mushroom spines are increased during sleep following re-exposure to fear conditioning 28 days after the initial exposure. (D) Volume and head diameter of Arc+ mushroom spines is increased during sleep following re-exposure to fear conditioning, along with decreased neck length indicating strengthening of these synapses. Broad morphological changes also occur in Arc-spines during sleep suggesting more complex remodeling in these spines during sleep following re-exposure to a traumatic event .

### Engram specific upscaling vs broad downscaling in CA1 during sleep

Previous studies, including from our group, reported broad downscaling of dendritic spines in CA1 during sleep ^40,51^, opposite to our current findings of selective upscaling of dendritic spines following fear learning. The broad downscaling in our previous study occurred in mice without recent learning ^40^, and the study in adolescent mice was conducted following exposure to novel objects ^51^, which does not represent the biologically relevant emotional learning that occurs during fear conditioning. Broad downscaling during sleep thus may be a standard process to enhance signal to noise ratio following standard contextual encounters, whereas recent emotional learning from that occurs during stressful experiences that may be necessary for survival results in selective upscaling during sleep in the same hippocampal region.

Recent evidence demonstrated that sleep following contextual fear conditioning in mice suppressed cFos and Arc protein expression in the CA1 region compared to mice that underwent 6 hours of SD ^52^. Furthermore, sleep following contextual fear conditioning also suppressed phosphorylated S6 and ERK1/2 compared to mice that underwent SD, along with CA1 specific transcriptional alterations ^52^. Phosphorylated S6 and ERK1/2 are increased by neuronal activity and involved in promoting formation and maintenance of dendritic spines ^53–56^, suggesting these molecular processes and the reported associated transcriptional pathways may underlie our observed dendritic spine changes in Arc+ neurons following contextual fear learning.

### CA1 stratum oriens selective contextual fear memory engram changes during sleep

The majority of Arc+ dendrites observed in CA1 were located in the stratum radiatum (Fig. 2A). The CA1 stratum radiatum has been implicated as a key site of contextual fear memory consolidation ^57^. Apical dendrites of CA1 pyramidal neurons receive synaptic inputs from CA3 Schaffer collaterals onto mid and proximal apical segments ^58^, and are involved in the formation of emotional memory ^59,60^. Apical dendrites in CA1 contextual fear memory engram cells display increased density and size of dendritic spines receiving inputs from CA3 contextual fear memory engram cells following contextual fear learning ^61^. Our evidence for synaptic upscaling during sleep in Arc+ branches, including greater densities of mushroom spines and enhanced spine morphology (Fig. 2,4) suggests that connections from CA3 to CA1 following fear learning are strengthened during sleep. The spine morphology changes we observed, indicating enhanced memory storage during sleep in Arc+ spines immediately after fear learning, is in line with reports that spine morphology is enhanced in fear memory engram cells and prior studies suggest this correlates with fear memory strength ^62^. Furthermore, our findings indicate that SD after initial fear learning impairs strengthening of dendritic spines that occurs in engram cells as well as enlargement of the engram following re-exposure 28 days later (Fig. 5). Thus, SD after the initial fear learning may alter morphological and molecular processes that impact CA3 to CA1 connections that guide short and long-term memory consolidation and reconsolidation processes.

### Dendritic Spine morphological alterations

The volume and surface area of Arc+ thin spines was increased during sleep following initial fear learning, compared to the SD animals (Fig. 4). Greater surface area and volume indicates synaptic upscaling during sleep, as greater spine volume and surface area is associated with synaptic strength and density of glutamate receptors ^47–50^. Taken together with increased density of mushroom spines in Arc+ branches, this suggests that thin spines that participated in encoding contextual fear memory are in the process of synaptic enhancement during sleep and may develop into mushroom spines ^63^. Previous studies have demonstrated that increased synaptic plasticity is associated with a morphological shift from thin to mushroom spines ^64–67^. Arc-neurons in contrast did not display altered spine densities. Furthermore, the morphological changes observed including increased head and neck length of mushroom spines and stubby spines (Fig. 3), and decreased head and neck diameter, volume, and surface area of stubby spines (Fig. 4), indicate moderate downscaling of spines that are not involved in encoding contextual fear memory.

In comparison, fear re-exposure resulted in shorter neck length of mushroom spines in Arc+ branches during sleep (Fig. 6), which is associated with faster synaptic transmission and greater synaptic strength^44^. Furthermore, mushroom spines in Arc+ branches displayed greater head diameter and volume during sleep (Fig. 7), indicating morphological strengthening of mushroom spines during sleep following fear re-exposure in the original contextual fear engram. Arc-branches displayed shorter head and neck length of thin and mushroom spines, along with smaller volume and surface area of all spine types during sleep following re-exposure (Fig. 7). When taken together with increased density during sleep in Arc-branches (Fig. 5), this suggests more moderate morphological changes during sleep as neurons that were not part of the original contextual fear memory trace are recruited into this trace following re-exposure. Overall, these findings suggest that sleep after re-exposure to a traumatic experience strengthens the contextual fear memory trace during sleep both in neurons that originally encoded this memory as well as neurons that were not part of the original memory engram, indicating expansion of the engram.

### Relevance to PTSD

Our findings suggest that sleep after a traumatic experience strengthens dendritic spines in the contextual fear memory engram. Our data, together with previous studies demonstrate that SD following fear conditioning impairs contextual fear memory ^68,69^, including work from our group ^70^, which suggests that SD after a traumatic experience may alleviate PTSD risk by preventing the strengthening of dendritic spines that encode contextual fear memory. Evidence that sleep deprivation reduces emotional memories in human subjects ^21^ provides further support for this protective effect.

Multiple traumatic experiences increase the risk of developing PTSD as well as the severity of PTSD symptoms ^23–27^. Therefore, we tested the hypothesis that SD following the initial traumatic event (initial fear conditioning session) weakens synaptic upscaling in the contextual fear memory engram following re-exposure to the traumatic event 28 days later. Our findings demonstrate that in control mice, re-exposure to fear conditioning results in synaptic upscaling in the original contextual fear memory engram neurons as well as in neurons that were not part of the original engram, suggesting expansion of the contextual fear memory engram following re-exposure. Synaptic upscaling in neurons that were not part of the original memory trace may contribute to generalized contextual fear inherent in PTSD ^71,72^. Furthermore, SD following the initial fear conditioning session significantly decreased the upscaling that occurred following re-exposure, particularly in neurons that were not part of the original engram, and weakened freezing during re-exposure (Fig. 5). This suggests that SD following a traumatic experience may be a protective factor for potential re-exposure to a traumatic event and possibly alleviate PTSD risk and severity. Evidence that sleep after learning enhances emotional memories in humans for up to 4 years ^73^, provides support for such long-term protective effects of SD following emotional learning.

### Limitations

The lack of dendritic branch and segment specific analysis in our study is an important limitation. The limited distribution of Arc+ and Arc-neurons and their dendritic branches in CA1 concentrated in the virus injection site made it challenging to follow branches from individual neurons. However, obtaining information regarding dendritic spines measures from neurons that were part of the contextual fear memory engram vs neurons that were not in SD and control animals for 3-dimensional high resolution dendritic spine quantification was important. Therefore, we analyzed all clearly visible Arc+ and Arc-labeled dendritic branches in our samples. Lack of cell type specificity for the dendritic spines sampled is another potential limiting factor in understanding how dendritic spines may be altered during sleep in specific populations of excitatory and inhibitory neurons. Arc is only expressed in excitatory neurons ^74^, thus our design using ArcCreER^T2^ mice allowed for selective analysis of dendritic spines in excitatory neurons. Future studies designed to examine dendritic spines in inhibitory neurons may reveal differential dendritic spine changes during sleep in inhibitory neuron populations, as suggested by recent evidence for a key role of hippocampal somatostatin neurons during sleep ^75^. Hippocampal region-specific molecular alterations have been reported in SD mice following contextual fear conditioning ^52^. This suggests dendritic spine changes may vary by region. Future studies will examine engram specific dendritic spine changes in SD mice from other hippocampal sectors.

## Conclusion

In summary, our findings represent the first evidence for contextual fear memory engram dendritic spine changes in the hippocampus following SD. Collectively, our data indicate upscaling of dendritic spines during sleep in the contextual fear memory engram. Furthermore, our findings suggest that later re-exposure to contextual fear conditioning expands the fear memory engram and indicate that SD following initial contextual fear learning impairs synaptic strengthening during sleep that occurs following future re-exposure to a traumatic experience.

## Data Availability

The data and original contributions presented in this study are included in the manuscript and are available upon request to the corresponding author.

## Acknowledgments

This work was funded by support from NIMH R21MH117460 and NIGMS P20GM144041.

